## Supplemental material for "Deep learning-driven fragment ion series classification enables highly precise and sensitive de novo peptide sequencing"

**- Supplementary Information**

Daniela Klaproth-Andrade[1], Johannes Hingerl[1], Nicholas H. Smith[1], Jakob Träuble[1], Mathias Wilhelm[2,*], Julien Gagneur[1,3,4,5,*]

[1] Computational Molecular Medicine, School of Computation, Information and Technology, Technical University of Munich, Munich, Germany
[2] Computational Mass Spectrometry, School of Life Sciences, Technical University of Munich, Munich, Germany
[3] Institute of Human Genetics, School of Medicine, Technical University of Munich, Munich, Germany
[4] Computational Health Center, Helmholtz Center Munich, Neuherberg, Germany
[5] Munich Data Science Institute, Technical University of Munich, Garching, Germany
* Corresponding authors

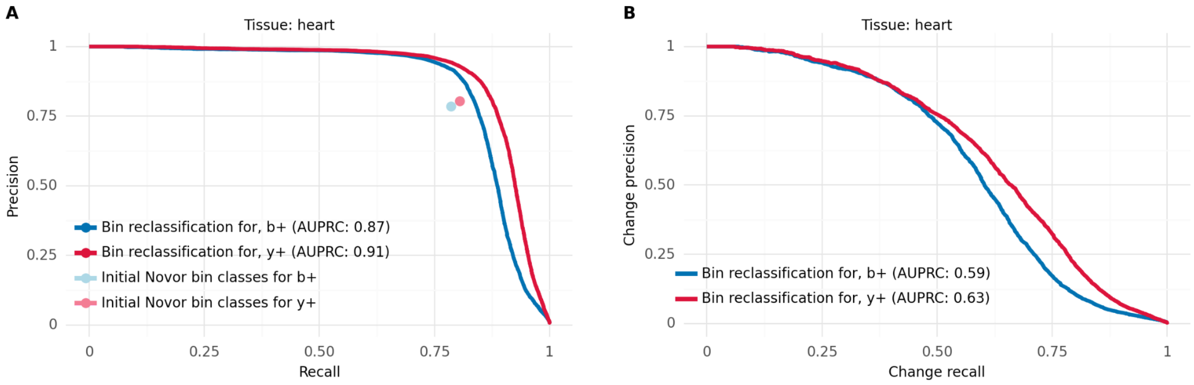


**Supplementary Figure 1 | Bin reclassification performance for Novor-predicted peptides on heart. A,** Precision-recall curves for bin reclassification of b+ and y+ ion series after relabeling initial bin classes proposed by Novor on the test set of the heart sample compared to the precision and recall computed at bin level for the initial class labeling. **C,** Change-precision-recall curves at bin level for b+ and y+ ions on the test set of the heart sample after relabeling initial bin classes proposed by Novor on the heart sample.

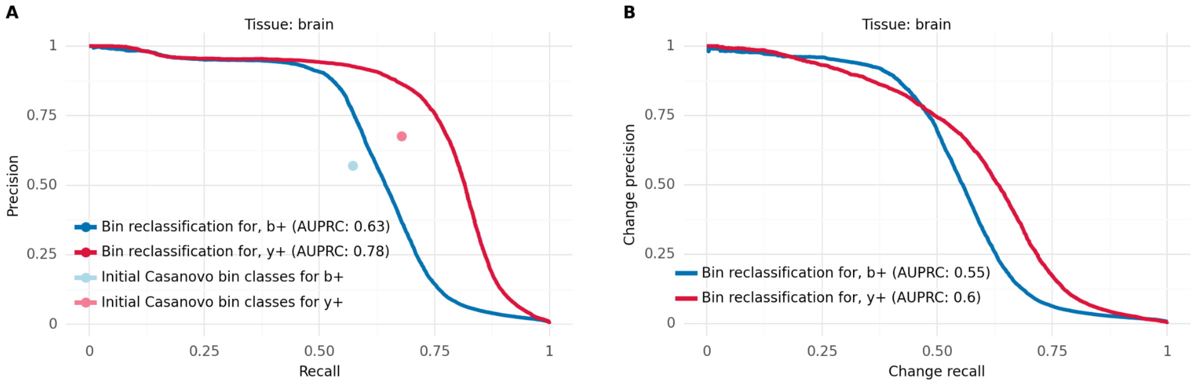


**Supplementary Figure 2 | Bin reclassification performance for Casanovo-predicted peptides on brain. A,** Precision-recall curves for bin reclassification of b+ and y+ ion series after relabeling initial bin classes proposed by Casanovo on the test set of the brain sample compared to the precision and recall computed at bin level for the initial class labeling. **C,** Change-precision-recall curves at bin level for b+ and y+ ions on the test set of the heart sample after relabeling initial bin classes proposed by Casanovo on the brain sample.

**Supplementary Table 1 | Feature list and feature importances.** Complete list of 114 features used as input for the random forest regressor to estimate the Levenshtein distance of a peptide candidate to the correct peptide sequence. The feature number indicates the order in which the features are passed to the model. The feature category indicates one of the three possible categories: similarity, counting, and bin reclassification-based features. The subcategory indicates which metric was used for feature computation or described how the feature was computed. The specification indicates whether the feature was computed on all peaks or bins or whether only a subset was considered (only b- or y-ions or only bins with predicted change probability above a certain threshold) or whether a logarithmic transformation (log2) was applied prior to feature computatio on test set.

| Category | Subcategory | Feature number | Specification | Importance |
|---|---|---|---|---|
| Similarity between matching experimental and PROSIT-predicted peaks | Spectral angle | 1 | all ions | 0.0071 |
| | | 2 | b ions | 0.0017 |
| | | 3 | y ions | 0.0010 |
| | Pearson correlation coefficient | 4 | all ions | 0.0018 |
| | | 5 | b ions | 0.0021 |
| | | 6 | y ions | 0.0016 |
| | | 7 | all ions with log2 normalization | 0.0158 |
| | | 8 | b ions with log2 normalization | 0.0016 |
| | | 9 | y ions with log 2 normalization | 0.0059 |
| | Cosine similarity | 10 | all ions | 0.0054 |
| | | 11 | b ions | 0.0015 |
| | | 12 | y ions | 0.0039 |
| | | 13 | all ions with log2 normalization | 0.0045 |
| | | 14 | b ions with log2 normalization | 0.0010 |
| | | 15 | y ions with log 2 normalization | 0.0020 |
| | Mean of absolute peak differences | 16 | all ions | 0.0269 |
| | | 25 | b ions | 0.0017 |
| | | 34 | y ions | 0.0012 |
| | | 43 | all ions with log2 normalization | 0.0018 |
| | | 52 | b ions with log2 normalization | 0.0011 |
| | | 61 | y ions with log 2 normalization | 0.0016 |
| | Standard deviation of absolute peak differences | 17 | all ions | 0.0030 |
| | | 26 | b ions | 0.0018 |
| | | 35 | y ions | 0.0017 |
| | | 44 | all ions with log2 normalization | 0.0013 |

| | | | | |
|---|---|---|---|---|
| | | 53 | b ions with log2 normalization | 0.0010 |
| | | 62 | y ions with log 2 normalization | 0.0024 |
| | 3rd quartile of absolute peak differences | 18 | all ions | 0.0010 |
| | | 27 | b ions | 0.0017 |
| | | 36 | y ions | 0.0055 |
| | | 45 | all ions with log2 normalization | 0.0014 |
| | | 54 | b ions with log2 normalization | 0.0015 |
| | | 63 | y ions with log 2 normalization | 0.0233 |
| | 2nd quartile of absolute peak differences | 19 | all ions | 0.0008 |
| | | 28 | b ions | 0.0017 |
| | | 37 | y ions | 0.0015 |
| | | 46 | all ions with log2 normalization | 0.0016 |
| | | 55 | b ions with log2 normalization | 0.0011 |
| | | 64 | y ions with log 2 normalization | 0.0105 |
| | 1st quartile of absolute peak differences | 20 | all ions | 0.0007 |
| | | 29 | b ions | 0.0015 |
| | | 38 | y ions | 0.0020 |
| | | 47 | all ions with log2 normalization | 0.0012 |
| | | 56 | b ions with log2 normalization | 0.0010 |
| | | 65 | y ions with log 2 normalization | 0.0027 |
| | Minimum of absolute peak differences | 21 | all ions | 0.0039 |
| | | 30 | b ions | 0.0013 |
| | | 39 | y ions | 0.0033 |
| | | 48 | all ions with log2 normalization | 0.0016 |
| | | 57 | b ions with log2 normalization | 0.0014 |
| | | 66 | y ions with log 2 normalization | 0.0088 |
| | Maximum of absolute peak differences | 22 | all ions | 0.0012 |
| | | 31 | b ions | 0.0013 |
| | | 40 | y ions | 0.0011 |
| | | 49 | all ions with log2 normalization | 0.0025 |
| | | 58 | b ions with log2 normalization | 0.0016 |
| | | 67 | y ions with log 2 normalization | 0.0171 |
| | MSE of peak differences | 23 | all ions | 0.0018 |
| | | 32 | b ions | 0.0018 |

| | | | | |
|---|---|---|---|---|
| | | 41 | y ions | 0.0012 |
| | | 50 | all ions with log2 normalization | 0.0012 |
| | | 59 | b ions with log2 normalization | 0.0017 |
| | | 68 | y ions with log 2 normalization | 0.0077 |
| | Dot product ob peak differences | 24 | all ions | 0.0081 |
| | | 33 | b ions | 0.0015 |
| | | 42 | y ions | 0.0013 |
| | | 51 | all ions with log2 normalization | 0.0104 |
| | | 60 | b ions with log2 normalization | 0.0020 |
| | | 69 | y ions with log 2 normalization | 0.0045 |
| | Spearman correlation coefficient | 70 | all ions | 0.3163 |
| | | 71 | b ions | 0.0010 |
| | | 72 | y ions | 0.0013 |
| | | 73 | all ions with log2 normalization | 0.3033 |
| | | 74 | b ions with log2 normalization | 0.0010 |
| | | 75 | y ions with log 2 normalization | 0.0013 |

| | | | | |
|---|---|---|---|---|
| Counting between matching experimental and PROSIT-predicted peaks | Number of peaks with non-zero experimental intensity and non-zero PROSIT-intensity | 76 | all ions | 0.0036 |
| | | 77 | b ions | 0.0033 |
| | | 78 | y ions | 0.0036 |
| | Number of peaks with non-zero experimental intensity and non-zero PROSIT-intensity relative to total amount of PROSIT-intensities | 79 | all ions | 0.1397 |
| | | 80 | b ions | 0.0087 |
| | | 81 | y ions | 0.0081 |
| | Number of peaks with non-zero experimental intensity and non-zero PROSIT-intensity relative to total amount of non-zero PROSIT-intensities | 82 | all ions | 0.0101 |
| | | 83 | b ions | 0.0021 |
| | | 84 | y ions | 0.0003 |
| | Number of peaks with non-zero experimental intensity and zero PROSIT-intensity | 85 | all ions | 0.0039 |
| | | 86 | b ions | 0.0006 |
| | | 87 | y ions | 0.0051 |
| | Number of peaks with non-zero experimental intensity and zero PROSIT-intensity relative to total amount of PROSIT-intensities | 88 | all ions | 0.0054 |
| | | 89 | b ions | 0.0019 |
| | | 90 | y ions | 0.0061 |
| | Number of peaks with non-zero | 91 | all ions | 0.0052 |

| | | | | |
|---|---|---|---|---|
| | experimental intensity and zero PROSIT-intensity relative to total amount of non-zero PROSIT-intensities | 92 | b ions | 0.0012 |
| | | 93 | y ions | 0.0054 |
| | Number of peaks with zero experimental intensity and non-zero PROSIT-intensity | 94 | all ions | 0.0434 |
| | | 95 | b ions | 0.0014 |
| | | 96 | y ions | 0.0035 |
| | Number of peaks with zero experimental intensity and non-zero PROSIT-intensity relative to total amount of PROSIT-intensities | 97 | all ions | 0.0067 |
| | | 98 | b ions | 0.0011 |
| | | 99 | y ions | 0.0082 |
| | Number of peaks with zero experimental intensity and non-zero PROSIT-intensity relative to total amount of non-zero PROSIT-intensities | 100 | all ions | 0.0179 |
| | | 101 | b ions | 0.0023 |
| | | 102 | y ions | 0.0003 |
| | Number of peaks with non-zero experimental intensity | 103 | all ions | 0.0060 |
| | | 104 | b ions | 0.0016 |
| | | 105 | y ions | 0.0121 |
| | Number of peaks with non-zero experimental intensity relative to total amount of PROSIT-intensities | 106 | all ions | 0.0039 |
| | | 107 | b ions | 0.0031 |
| | | 108 | y ions | 0.0035 |
| Binreclassification-based | Number of proposed bin changes above change probability threshold | 109 | threshold: 0.3 | 0.0335 |
| | | 110 | threshold: 0.4 | 0.0559 |
| | | 111 | threshold: 0.45 | 0.0828 |
| | | 112 | threshold: 0.5 | 0.1145 |
| | | 113 | threshold: 0.55 | 0.0795 |
| | | 114 | threshold: 0.6 | 0.1266 |

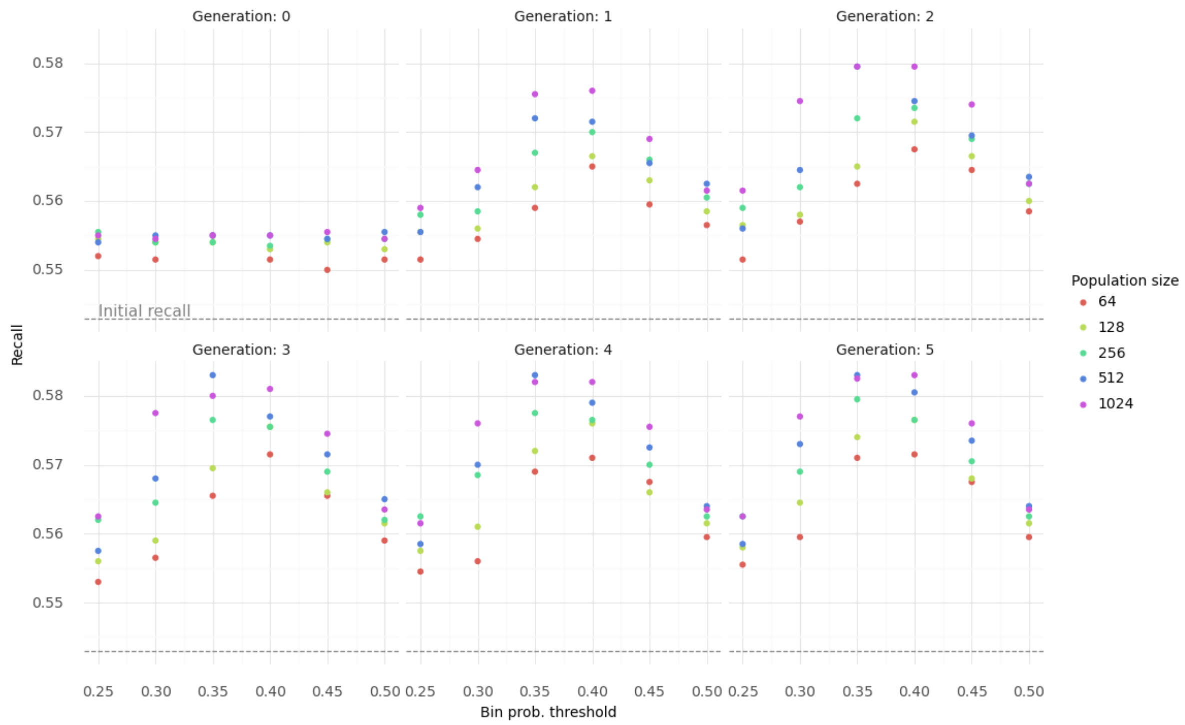


**Supplementary Figure 3 | Performance comparison with different hyper-parameters.** Peptide recall after running Spectralis with different selections of bin probability threshold, number of generations and population size on a subset of the heart sample.

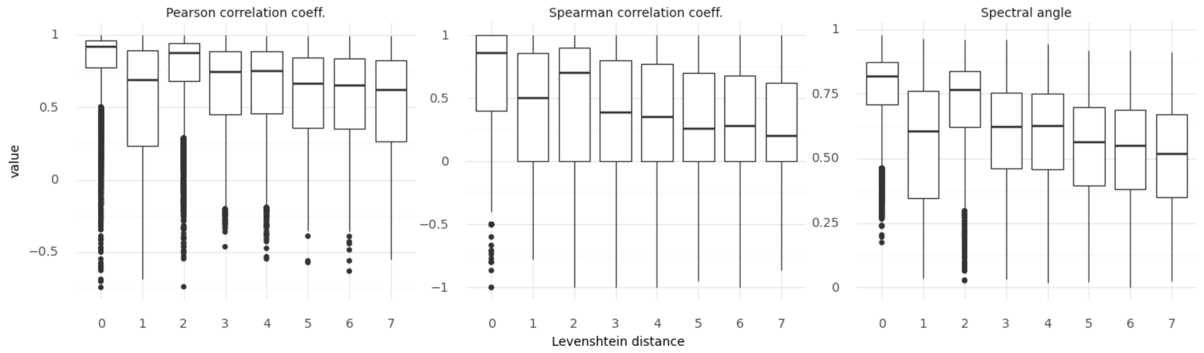


**Supplementary Figure 4 | Distribution of features against different Levenshtein distances.** Pearson and Spearman correlation coefficients, as well as spectral angles against Levenshtein distance of peptide identifications by Novor and Casanovo to the correct peptide sequence by MaxQuant across all 30 human samples for initial Levenshtein distances smaller than 8.

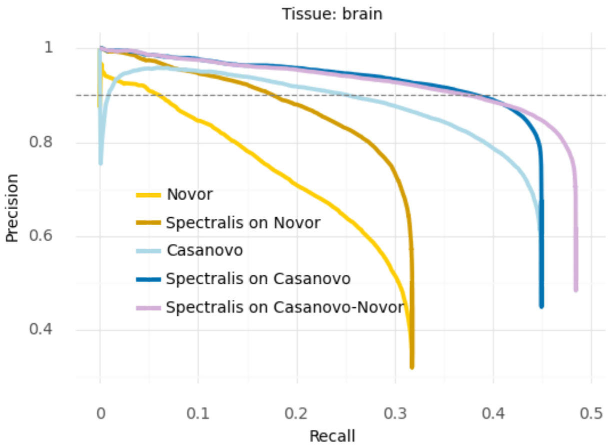


**Supplementary Figure 5 | Levenshtein distance estimator performance on the brain sample.** Precision-recall curves at peptide level before and after rescoring peptide identifications by Novor and Casanovo on the brain sample with the trained regression model that estimates Levenshtein distances, including the precision and recall for the combination of Casanovo and Novor sequences (Casanovo-Novor).

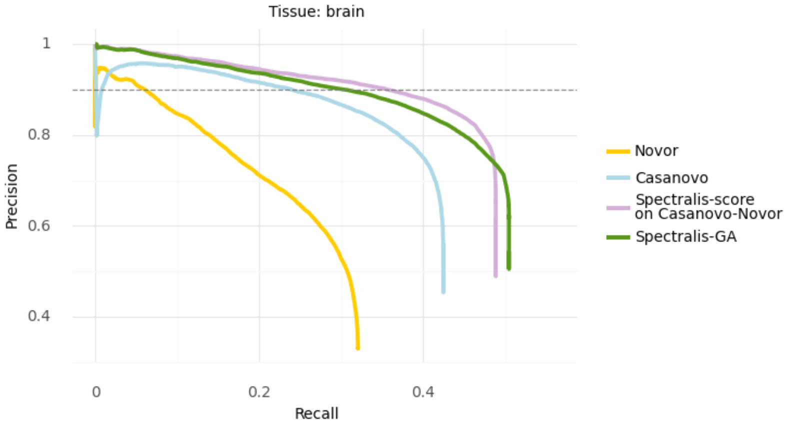


**Supplementary Figure 6 | Spectralis performance on the brain sample.** Precision-recall curves of identifications at peptide level for Novor, Casanovo, and Spectralis on the test set of the brain sample including the precision and recall for the combination of Casanovo and Novor sequences (Casanovo-Novor).

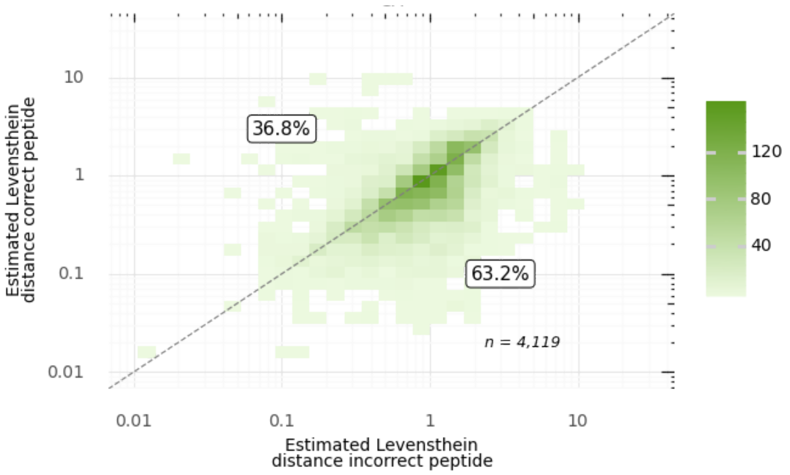


**Supplementary Figure 7 | Comparison of estimated Levenshtein distances for correct and incorrect peptides with a Levenshtein distance of two.** Estimated Levenshtein distances of incorrect peptides against the estimated Levenshtein distances of the corresponding correct peptide sequences for a given spectrum across all 30 samples for peptide sequences with an actual Levenshtein distance of two to the correct peptide.

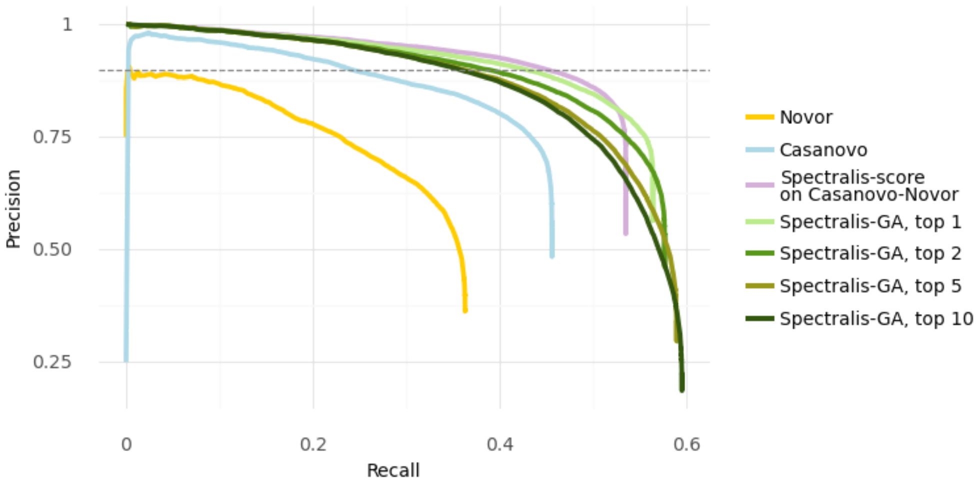


**Supplementary Figure 8 | Spectralis performance on the heart sample including multiple peptide predictions per spectrum.** Precision-recall curves of identifications at peptide level for Novor, Casanovo and Spectralis-GA on the test set of the heart sample, including the precision and recall for the combination of Casanovo and Novor sequences (Casanovo-Novor) and the Spectralis-GA returning the top 1,2,5 and 10 best scored peptide candidates per spectrum.

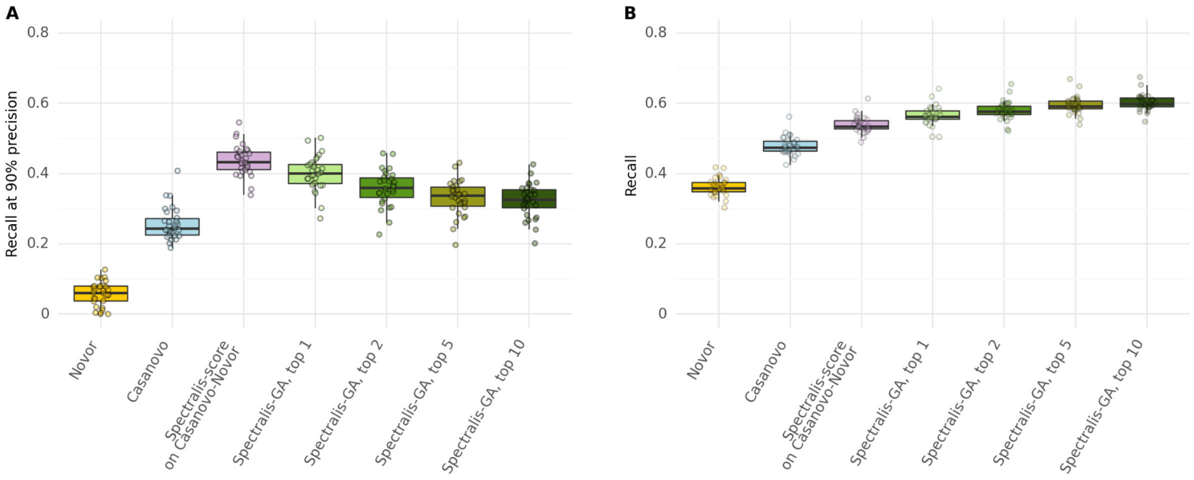


**Supplementary Figure 9 | Spectralis performance on all samples including multiple peptide predictions per spectrum. A,** Recall at 90% precision for Novor, Casanovo, Spectralis-score on the combination of Casanovo and Novor (Casanovo-Novor) and Spectralis-GA returning the top 1,2,5 and 10 best score peptide candidates on the test sets of all 30 samples. **E,** Overall recall for Novor, Casanovo, Spectralis-score Casanovo-Novor, and Spectralis-GA returning the top 1,2,5 and 10 best score peptide candidates on the test sets of all 30 samples.

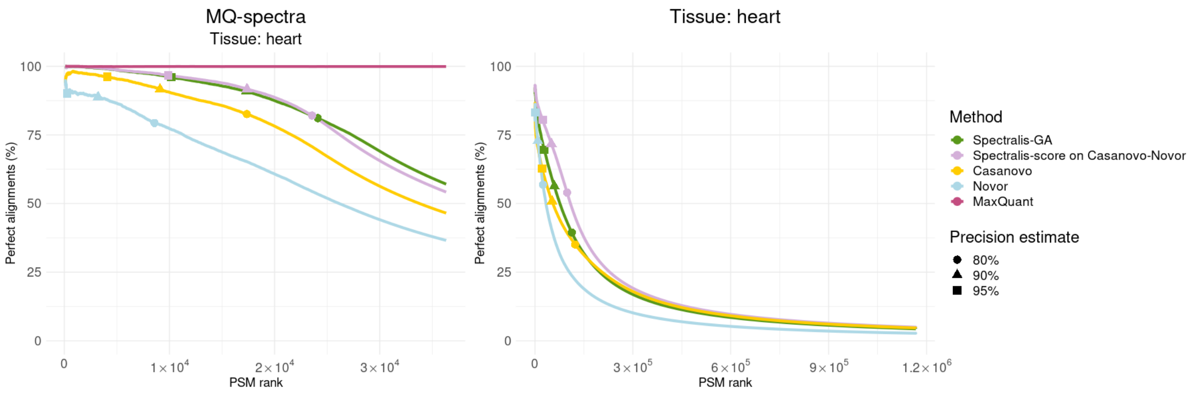


**Supplementary Figure 10 | Percentage of perfect alignments for heart sample.** Left: percentage of perfect alignments queried on peptide sequences queried on peptide sequences by MaxQuant, Novor, Casanovo and Spectralis-score on Casanovo-Novor and Spectralis-GA against a set of known and predicted gene translations using blastp on the set of spectra identified by MaxQuant (MQ-spectra, $n$ = 36,312) of the heart sample. For clarity, the first 100 peptides are omitted. Right: same as left, but for spectra not identified by MaxQuant (non-MQ-spectra, $n$ = 1,167,029).

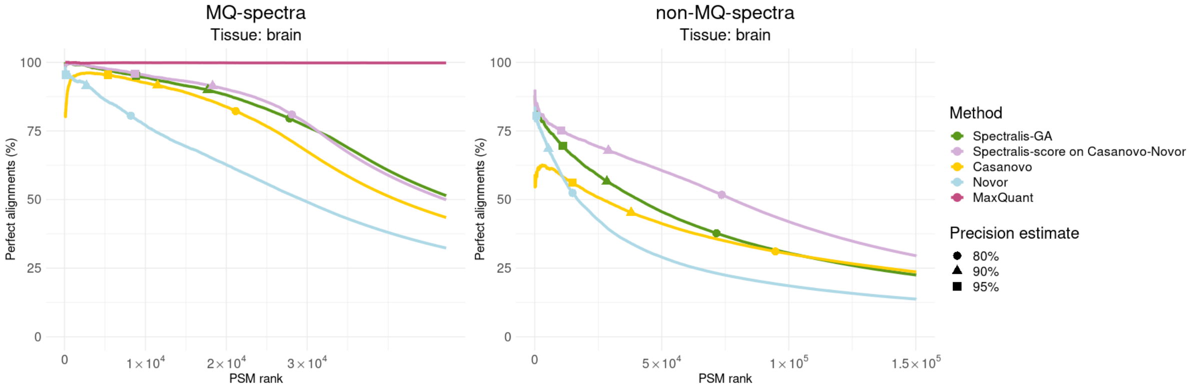


**Supplementary Figure 11 | Percentage of perfect blastp matches for brain sample.** Left: Percentage of perfect alignments queried on peptide sequences queried on peptide sequences by MaxQuant, Novor, Casanovo and Spectralis-score on Casanovo-Novor and Spectralis-GA against a set of known and predicted gene translations using blastp on the set of spectra identified by MaxQuant (MQ-spectra, $n$ = 47,181) of the brain sample. For clarity, the first 100 peptides are omitted. Right: same as left, but for spectra not identified by MaxQuant (non-MQ-spectra) showing only the top 150,000 ranked peptide candidates of each method (out of 1,040,253).
